# Nanoencapsulation of clove oil and study of physico-chemical properties, cytotoxic, haemolytic and antioxidant activities

**DOI:** 10.1101/2020.08.28.269886

**Authors:** Pramod G Nagaraju, Parineeta Sengupta, C. G. Poornima Priyadarshini, Pooja J Rao

**Author notes:** To whom correspondence should be addressed: Dr.Pooja J Rao, Spice & Flavour Sciences, CSIR- Central Food Technological Research Institute, Mysuru- 570020, Dr. Poornima Priyadarshini C G, Department of Molecular Nutrition, Central Food Technological Research Institute, Mysuru- 570020.

## Abstract

The therapeutic properties of clove oil is known for centuries, however, the pungent nature, chemical instability and low water solubility impose limitations in harnessing its therapeutic potential. Hence, nanoencapsulation of clove oil was performed to overcome the above constraints and control its in-vitro release. The stability of nanoemulsion depends on various factors where the surfactant and its hydrophile/lipofile balance (HLB) play a key role. The non-ionic surfactants Tween 20, 40 and 80 with HLB of 16.7, 15.6 and 15, respectively, were used to study the stability of clove oil nanoemulsion (CON). The creaming index of CON prepared with Tween 20, 40 and 80 was 22.75 and 17.5 and 1.5%, respectively, after 8 days of storage at room temperature. Tween 20 and 40 produced particles > 300 nm while Tween 80 resulted in particles of size ∼150 nm. Transmission electron microscopic image of spray dried CON prepared with Tween 80 showed particle size in the range 150-190 nm after one month of storage at room temperature. The in vitro release studies showed 76% and 42% cumulative release of CON and native clove oil (NC), respectively at pH 7.4. The cellular toxicity of CON was significantly reduced by four fold compared to NC at a concentration of 60 µg/mL when tested on Caco2 cells. Similarly, haemolytic activity on red blood cells revealed less than 10% haemolysis signifying the compatibility of CON for its nutraceutical applications. In addition, CON also exhibited higher in-vitro antioxidant compared to NC as shown by DPPH and ABTS radical scavenging activity. Collectively, we have developed a unique method for NC nanoencapsulation using cost effective polysaccharide (maltodextrin) and surfactant for stabilizing the nanoemulsion for increased bioactivity.

## 1. Introduction

Clove oil, an essential oil extracted from the spice *Syzigum aromaticaum*, is known for its therapeutic properties such as antioxidant [1], anti-inflammatory [2], anti-aging [3] and activity against pathogenic organisms [4]. These activities are attributed to the presence of unsaturated phenolic compounds mainly eugenol, eugenyl acetate and β-caryophyllene (Fig. 1). Clove oil is used as a food additive at approved concentration of <1500 ppm while toxicity has been reported beyond a concentration of 3.75 g/kg body weight [1, 5]. Its use as a nutraceutical ingredient is limited due to low water solubility, poor stability in ambient conditions, high pungency, cytotoxicity and reduced bioavailability [6, 7]. The organoleptic characteristics change during conventional storage, as the components present in essential oils easily undergo oxidation, isomerization, cyclization or dehydrogenation reactions, triggered either due to enzymatic or chemical reaction [8]. Thus, to avoid clove oil degradation during storage and processing encapsulation in carrier materials such as lecithin, arabica gum-whey protein concentrate, poly-lactic glycolic acid were used. This also resulted in the enhancement of its water solubility [9, 10]. However, maltodextrin, widely used as food additive, has not been explored to prepare clove oil nanoemulsions.

**Figure 1.**
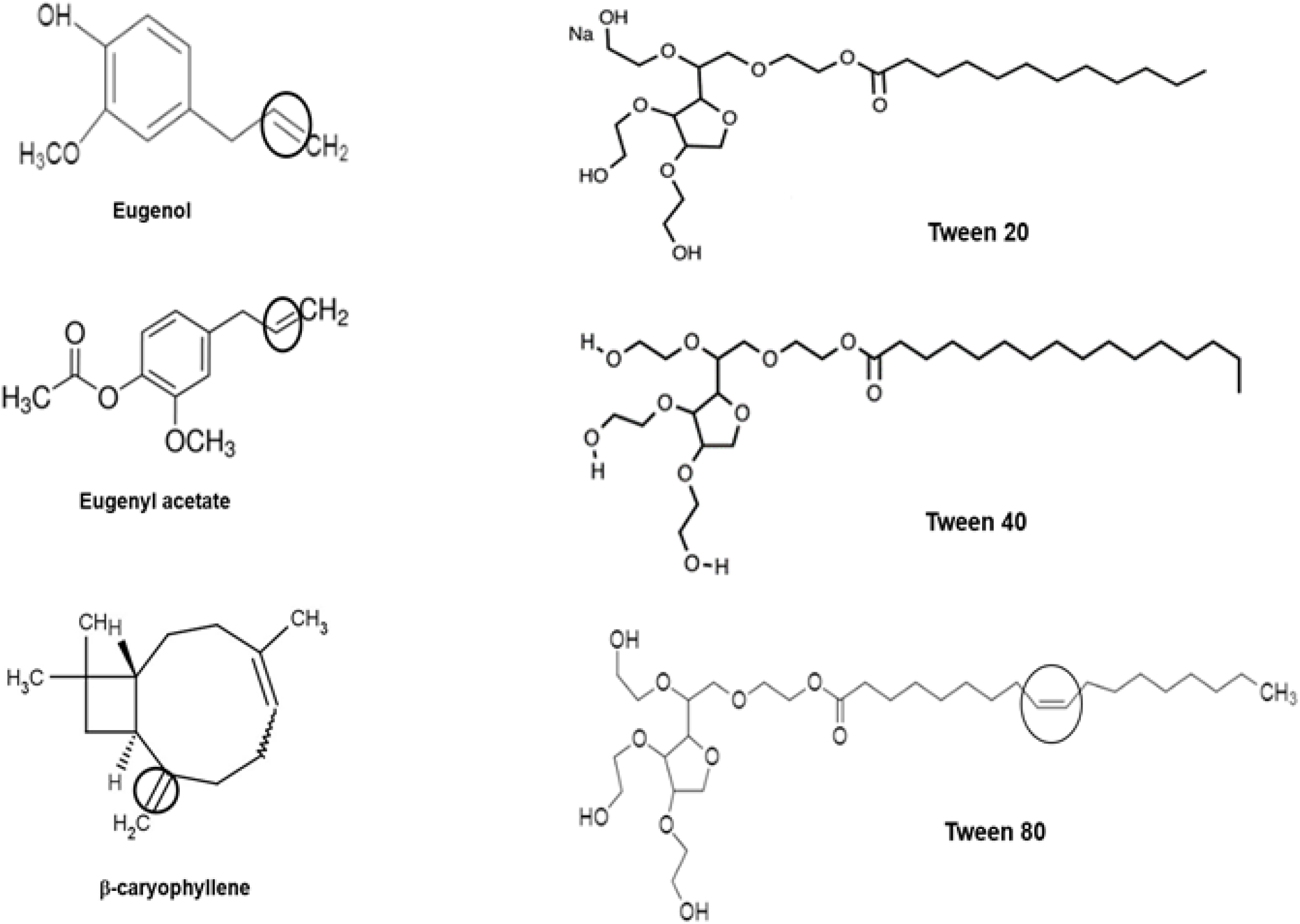
Chemical structure of clove oil components and surfactants used in the study. The unsaturated bond of Tween 80 and allyl group of clove oil bioactives involved in the interaction are encircled.

Maltodextrin is a hygroscopic modified polysaccharide with different dextrose equivalent (DE) 3-20. It is widely used because of its gel forming ability, low cost and easy availability. Maltodextrin with low DE (<10) possess better gel forming ability than the high DE value because of its long oligosaccharides chain [11]. The use of maltodextrin in emulsion formation is critical, as it has no surface-active sites to adsorb on the emulsion droplet and stabilize the emulsion. Hence, the use of emulsifier or surfactant is inevitable to achieve a long-term stability of nanoemulsion. Surfactant stabilizes the emulsion by inserting their non-polar tail in helical coil of maltodextrin to adsorb on the droplet and reduce the surface tension at the interface [12, 13]. However, the concentration and the nature of emulsifier play a crucial role in influencing the emulsion stability [14]. Thus, in the present study, polysorbate based different non-ionic surfactants such as Tween 20, 40 and 80 (Fig. 1) have been studied to stabilize the maltodextrin-based clove oil emulsion.

The anti-bacterial property of clove oil nanoemulsion [15] alone/in combination with cinnamon oil [16], anti-fungal properties of eugenol, clove oil nanoemulsion alone/in combination with other spices have been reported [17-19]. In addition to this, the antioxidant, insecticidal and antiviral properties were also extensively studied [20, 21]. However, scanty reports are available for cytotoxicity and haemolytic activity of clove oil nanoemulsions. The present work focus on the influence of physico-chemical parameter on the stability of clove oil nanoemulsion, particle size of nanoencapsulated powder and controlled in vitro release under gastro-intestinal pH. In addition, the data on cytotoxicity, haemolytic activity and potential antioxidant activity of nanoencapsulated clove oil has also been presented.

## 2. Results and discussion

Clove oil nanoemulsion and spray dried nanoparticle powder prepared using maltodextrin and Tween 20/40/80, were studied for physico-chemical properties and biological activities.

### 2.1. Particle size, polydispersity index (PDI) and zeta (ζ) potential

The particle size of CON stabilized using Tween 20/40/80 was shown in Fig. 2 (a). The average particle size of nanoemulsion prepared with Tween 20, 40 and 80 were 241.6, 237.5 and 168 nm, respectively. A decrease in particle size was observed with decrease in the HLB of the surfactant. Tween 80 with low HLB (15) facilitated the formation of small particles. It is relatively more lipophilic than Tween 20 and 40 thus its interaction with oil droplets was more than other surfactants. It restricted the particle growth effectively by creating repulsive interaction between oil droplets at the interfacial layer. The chemical and steric structure of the surfactants also influence the size and intensity of the droplets/particles [22]. The CON prepared with Tween 20 and 40 showed broad peaks with wide distribution of particles in the size range of 80-900 nm along with small height peaks in the range 3500-6500 nm (or 3.5-6.5 µm). The CON prepared with Tween 80 showed uniform distribution of particles from 80-200 nm range with a small population of particles around 500 nm. Tween 20 and 40 possess saturated laurate-chain with 12 carbon atoms and palmitate-chain with 16 carbon atoms, respectively. Whereas, Tween 80 contains an oleate-chain with 18 carbon atoms and one unsaturated bond. This bond was responsible for establishing an interaction with clove oil bioactives that also contained unsaturated bonds (encircled in Fig. 1). The long non-polar chain of Tween 80 adsorbed effectively around the oil droplets and reduced the surface tension at interfacial layer by forming a monomer in the aqueous medium. A protective coating of surfactant around oil droplets by adsorbing to the oil-water interface helps in avoiding the aggregation of particles and forms small particles [14]. This also explains low PDI in Tween 80 compared to Tween 20 and 40. Table 1 showed the variation in particle size, and PDI of CON with Tween 20, 40 and 80 for a period of 8 days. The particle size of spray-dried powder was 171.1 nm and observed to be same during the storage period of a month.

**Table 1.** Influence of Tween 20, 40 and 80 on the Size (nm), Zeta potential (mV) and Polydispersity index (PDI) of Clove oil nanoparticle was measured by Zetasizer NanoZS (Malvern Instruments, Malvern, UK) at 28°C. (All values expressed are mean ± SD where n = 3).

| Days | SIZE |  |  | Zeta potential |  |  | PDI |  |  |
| --- | --- | --- | --- | --- | --- | --- | --- | --- | --- |
|  | T.20 | T.40 | T.80 | T.20 | T.40 | T.80 | T.20 | T.40 | T.80 |
| Day 0 | 241±0.2 | 237.5±1 | 168.7±1 | -<br>18.5±0.3 | -<br>24.4±0.1 | -<br>25.7±0.2 | 0.275 | 0.333 | 0.270 |
| Day 2 | 245±0.6 | 241±0.7 | 171.3±3 | -<br>17.2±0.2 | -<br>24.5±0.2 | -<br>25.2±0.3 | 0.269 | 0.331 | 0.271 |
| Day 4 | 259±1.2 | 249±1.0 | 173±2 | -<br>16.4±0.4 | -<br>21.2±0.3 | -<br>24.2±0.2 | 0.254 | 0.321 | 0.269 |
| Day 6 | 271±0.8 | 254±1.1 | 174±1 | -<br>15.6±0.5 | -<br>18.5±0.1 | -<br>23.9±0.1 | 0.244 | 0.295 | 0.266 |
| Day 8 | 275±1.6 | 261±0.8 | 182.5±2 | -14.6±1 | -<br>17.1±0.3 | -<br>22.7±0.6 | 0.245 | 0.301 | 0.265 |

**Figure 2.**
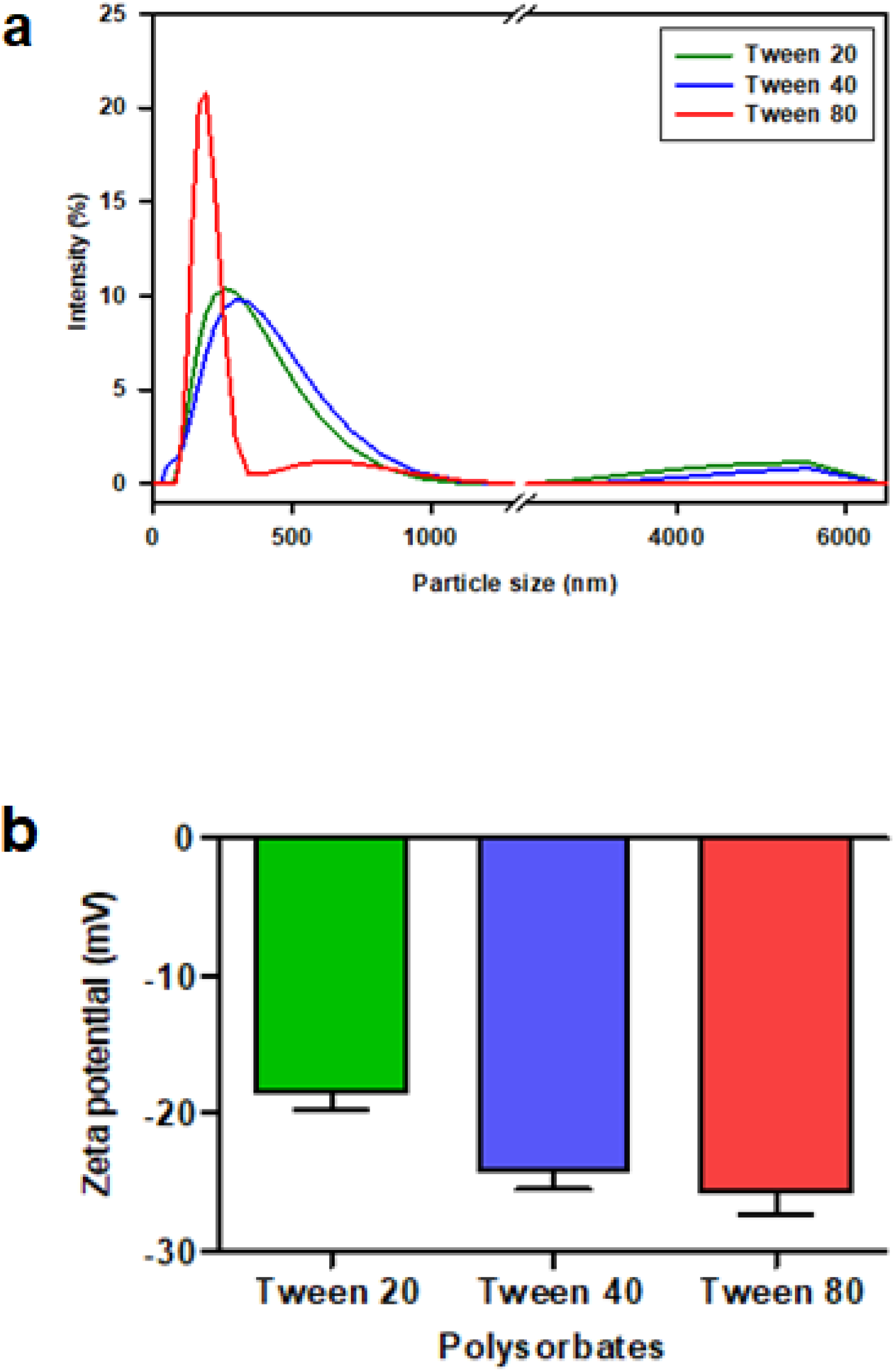
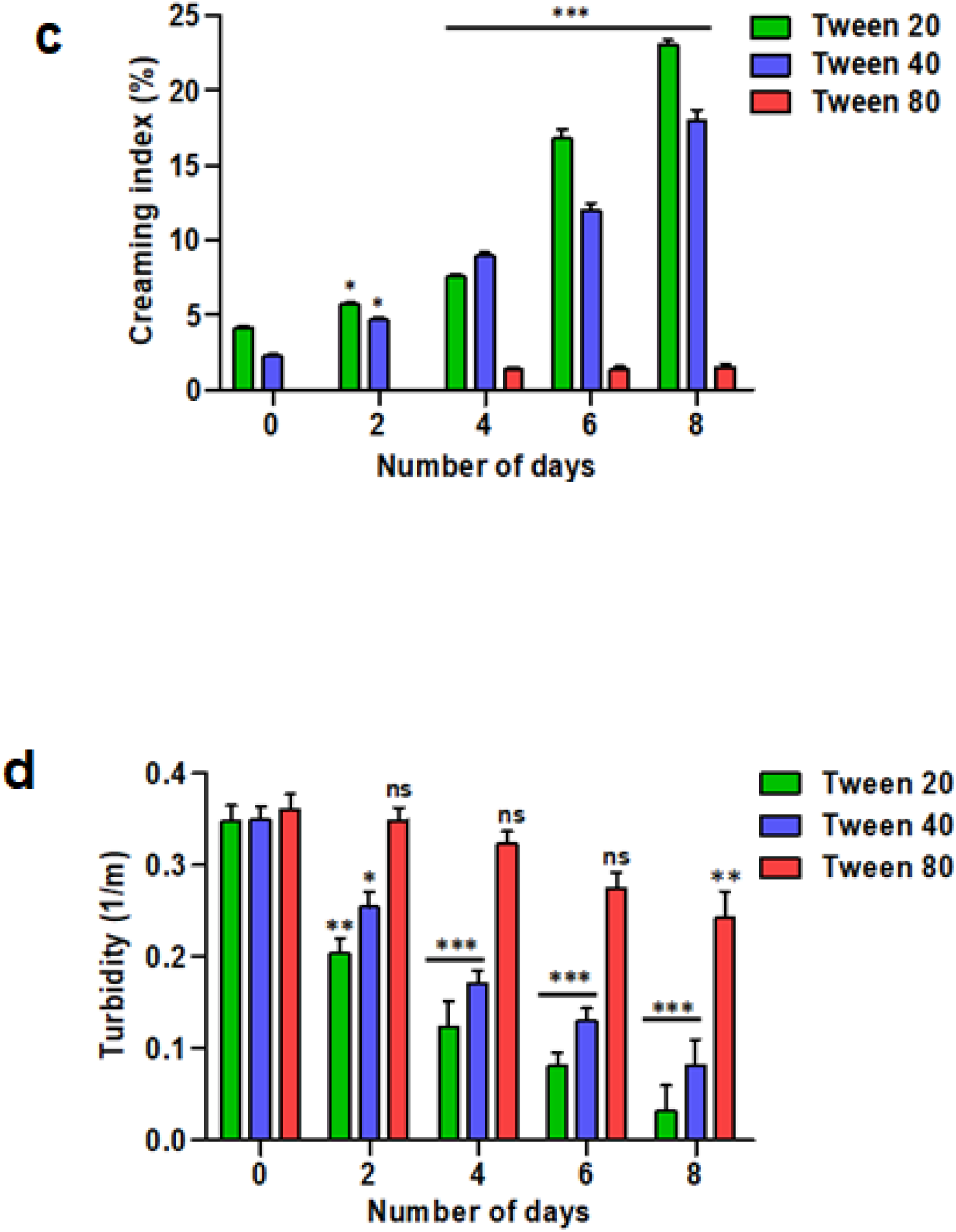
Influence of Tween 20, Tween 40 and Tween 80 on (a) Particle size (b) Zeta potential (c) Creaming index (d) Turbidity of Clove oil nanoparticle

The zeta potential value, an indicator of stability of nanoemulsion, showed an improvement with decrease in surfactant HLB (Fig. 2b). Tween 80 can interact effectively with the dispersed oil phase because of the presence of an unsaturated bond. Thus, it adsorbs strongly, stabilizes the emulsion and showed - 25.7 mV ζ value .The non-ionic surfactants stabilize the emulsion either sterically [23] or due to osmotic repulsive force [24] by forming a barrier around the oil droplet.

### 2.2. Emulsion stability

Appearance of a distinct oil layer (cream layer) is the initial stage of emulsion instability due to gravitational separation. The cream layer appeared in the CON samples prepared with Tween 20 and 40 on 0^th^ day itself indicated low stability of emulsions. The height of cream layer increased over the period and the creaming index was calculated as 22.75 and 17. 5 %, respectively (Fig. 2c). On the other hand, the CON with Tween 80 did not show any creaming until 4^th^ day and only few drops in immeasurable quantity were observed from 5^th^ day onwards. The calculated creaming index after 8^th^day was 1.5%. Phase separation was observed in all samples after 10-12 days. An increase in mean particle size over the period due to flocculation, coalescence lead to instability of droplets and finally gravitational/phase separation was observed [25].

The change/reduction in turbidity of emulsion is also an indicator of destabilizing nature of emulsion. The higher the scattering of light on emulsion droplets the higher the turbidity. It is clear from the figure (Fig. 2d) that the turbidity of CON samples with Tween 20, 40 linearly decreased. Whereas, CON with Tween 80 underwent changes after 4 days of storage. Turbidity of CON with Tween 20, 40 and 80 was calculated as 0.24, 0.03 and 0.08 m^-1^, respectively after 8 days of storage. The turbidity of the emulsion decreased because of three factors: (i) decrease in size of the droplet (ii) aggregation and settlement of droplets (iii) solubilization of oil droplets in surfactant micelles [23]. The sample CON with Tween-80 retain same particle-size, zeta potential with slight change in PDI properties over a period of 10 days thus, it was selected for further characterization.

### 2.3. Morphology

Fig. 3A and 3B shows the SEM and TEM images of maltodextrin and spray-dried CON. The native maltodextrin was crystalline and polymorphic in nature (Fig. 3A, a) whereas, CON spray dried powder was amorphous, spherical in shape with some dents (Fig. 3A, b-d). The CON powder prepared with Tween 20 and 40 show mostly large particles agglomerated with smaller ones (Fig. 3A, c). However, the CON prepared with Tween 80 showed several small and a few large individual particles (Fig. 3A, d) that could be due to the reduction in the steric interaction among particles. Figure 3B (a, b) shows the TEM images of maltodextrin and spray dried CON powders having particle size ∼2 µm and 150-200 nm, respectively.

**Figure 3.**
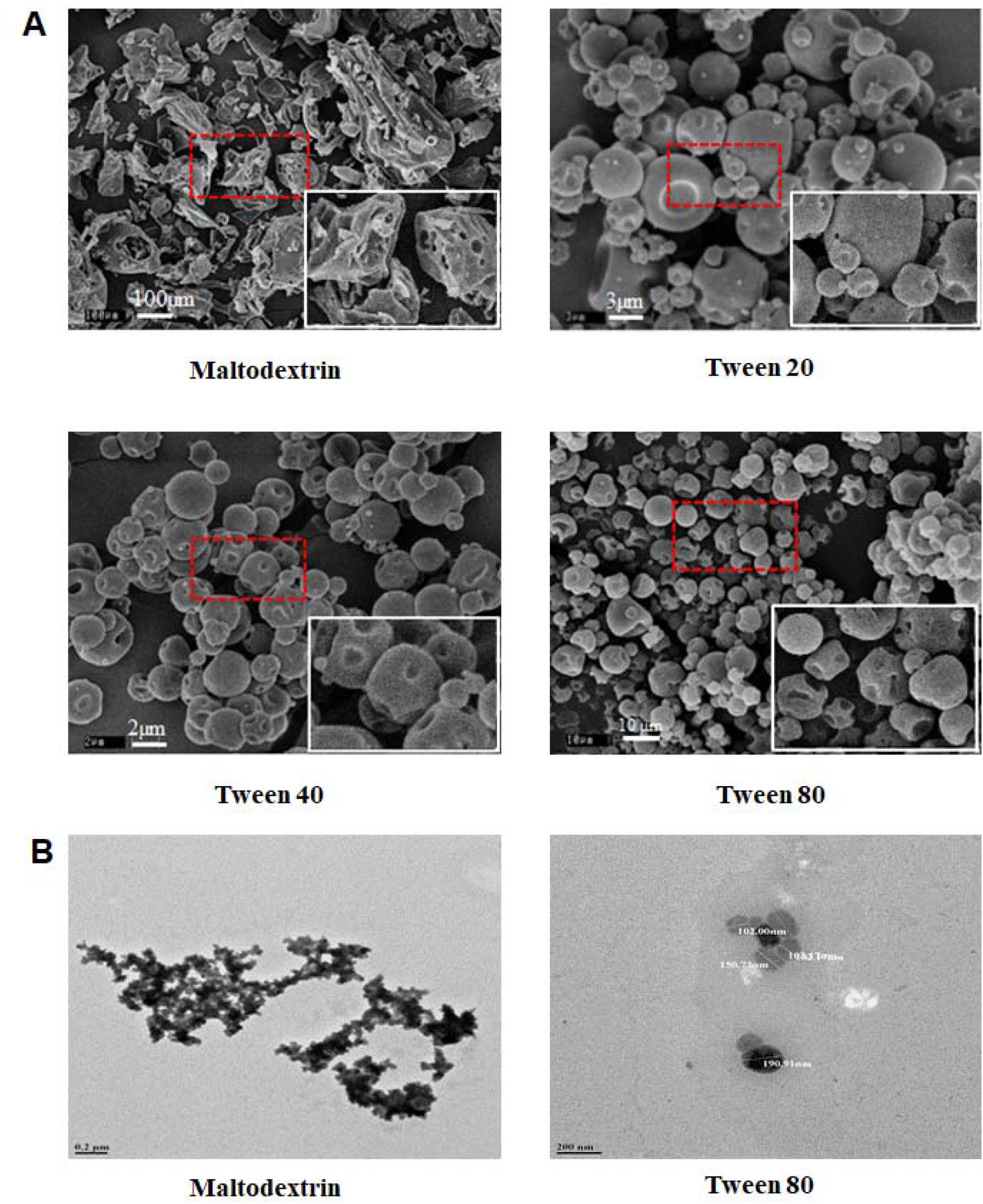
Morphology of clove oil nanoparticle (A) SEM images of Maltodextrin, Clove oil nanoparticle Spray-dried powder of Tween 20, 40, and 80 B) TEM of (a) Maltodextrin (b) Clove oil nanoparticle spray dried powder of Tween 80.

### 2.4. DSC, FTIR and GC

The DSC of NC, maltodextrin, and CON were performed to investigate their thermal stability (Fig. 4A). One endothermic peak at 204°C, indicating complete degradation, was observed in DSC thermogram of clove oil. The DSC profile of clove oil is in good agreement with the literature [26]. The kinetics of evaporation of essential oils was determined using both thermogravimetry and DSC profiles as described earlier [27]. The onset thermal evaporation temperature for clove oil and eugenol was 146°C and 162°C while the peak temperature was reported to be at 204°C and 209°C, respectively. DSC thermogram of maltodextrin did not show any peak up to 290°C, however, after 220°C heat absorption was more as indicated by a broad incomplete hump. Similar observations were reported by Elnaggar et al., 2010 where the authors used maltodextrin DE = 8-12 to develop oral disintegrating tablet under different preparation condition and found charring of sample after 220°C and no endothermic peak due to the melting of material [28]. In the present study, the CON and spray dried CON showed better stability compared to native clove oil. The DSC thermogram of CON did not show any peak until 290°C while the spray dried CON showed a small broad hump at 197°C. This may be due to the evaporation of surface adsorbed water and clove oil.

**Figure 4.**
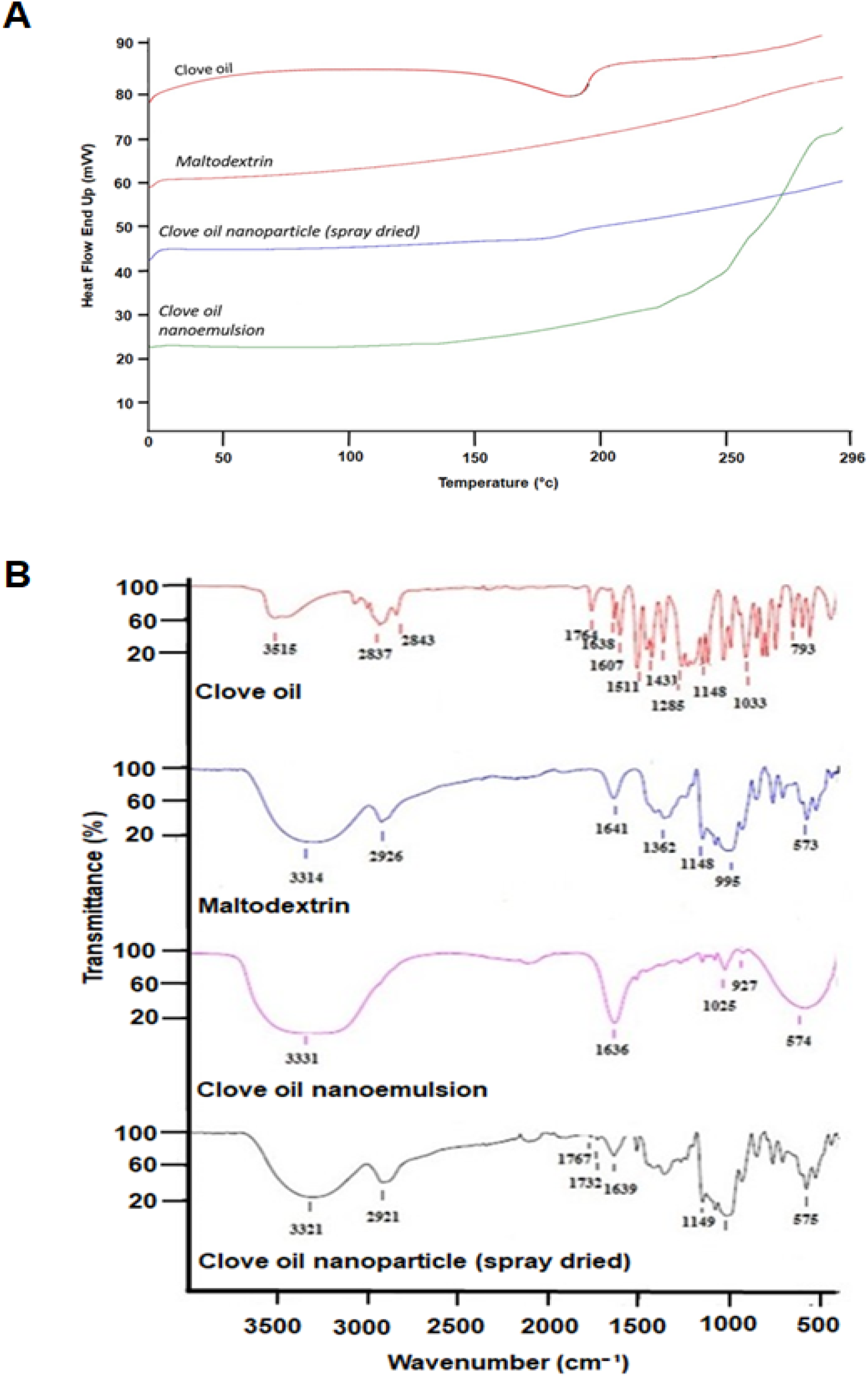
(A) Differential scanning calorimetry (DSC) profiles and (B) Fourier transform infrared spectroscopy (FTIR) of Clove oil, Maltodextrin, Clove oil nanoparticle.

Fig. 4B shows the FTIR peaks of NC, maltodextrin and CON. The Table 2 enlists the peaks observed due to bond vibration of functional groups and basic aromatic structure. The characteristic absorbance peaks of phenolic O−H and C−H stretching in aromatic ring at 3515 cm^-1^ [29] and 3076 cm^-1^, respectively were observed that were attributed to O−H group and aromatic structure of eugenol (a major compound in clove oil). The DC−H-stretching peak at 3003 cm^-1^ was observed due to allyl group (C−H attached to olefin) in eugenol [30]. Clove oil contains many substituted aromatic and sesquiterpene molecules and peaks responsible for aforementioned are listed in Table 2. Maltodextrin, a chain of D-glucose linked at α (1→4) glycosidic bond, mainly retained the FTIR peaks of basic molecule when it was present in emulsion and powder state of nanoencapsulated clove oil with few minor changes. The FTIR spectra of maltodextrin in native and emulsion form can be broadly divided in three regions. Peaks in the region extending 1500 to 600 cm^-1^ are due to vibration of COH, OCH, C−O, C−C bonds and the characteristic bands of open/pyranose ring structure of carbohydrate. A prominent band with small peaks ranging between 1150-930 cm^-1^ reflects the stretching vibration of C−O, C−C bonds. Peaks in the region 3310-3000 cm^-1^ are attributed to the vibration of O−H (symmetric stretching) and -C−H (asymmetric stretching) groups. The band in the region between 1770-1400 cm^-1^ and peaking at 1641 cm^-1^(native maltodextrin) is due to dCH_2_, dOCH, dCCH vibrations. In CON sample, the minor peaks of maltodextrin were merged and reduced while an enlargement of band between 1770-1400 peaking at 1639 cm^-1^ was observed. This could be due to the interaction between water molecule and maltodextrin leading for enhanced OC--H vibrations. Interestingly, spray dried maltodextrin encapsulating clove oil exhibits three small peaks at wavenumbers 1767 and 1732 cm^-1^. These peaks may be due to the -C=O stretch in acetate attached to aromatic ring indicating the presence of clove oil in the CON sample. It is understood that the peak responsible for O−H group of eugenol has probably submerged in the broad band of maltodextrin thus, not visible prominently.

**Table 2.** Assignment of FTIR peaks of Clove oil, Maltodextrin and Clove oil nanoparticle sample.

| Clove oil $\text{cm}^{-1}$ | Maltodextrin $\text{cm}^{-1}$ | CON emulsion/powder $\text{cm}^{-1}$ | Peak assignment |
| --- | --- | --- | --- |
| 3515 |  |  | O—H stretching in aromatic ring |
| 3076 |  |  | -C—H stretching in aromatic ring |
| 3003 |  |  | allyl group in eugenol |
| 2937, 2843 |  |  | C-H stretching aliphatic (could be from terpenes) |
| 1764 |  |  | -C=O stretch in acetate attached to aromatic ring |
| 1638 |  |  | C=C in allyl group |
| 1607, 1511, 1453 |  |  | C=C in aromatic ring |
| 1431 |  |  | CH<sub>2</sub> aliphatic |
| 1265-1200 |  |  | =C-O-C asymmetric stretch and =C-O-H from Ar-OH |
| 1033 |  |  | C-O-C stretching symmetric |
| 994, 911 |  |  | =C-H bending aliphatic |
| 793, 745 |  |  | C-H outplane bending |
|  | 1500 to 600 |  | vibration of COH, OCH, C—O, C—C bonds |
|  | 1150-930 |  | stretching vibration of C—O, C—C bonds |
|  | 3310-2900 |  | vibration of O—H (symmetric stretching) and -C—H(asymmetric stretching) groups |
|  | Band 1770 – 1400 peak 1641 |  | CH<sub>2</sub>+ OCH + CCH vibrations |
|  |  | 1767, 1739 | -C=O stretch in acetate attached to aromatic ring |

Fig. 5A shows the GC profile of NC and the clove oil extracted from the CON. It wa clear that encapsulation did not change the nature of clove oil, as the three peaks were prominently visible in both the figures. The presence of eugenol, eugenyl acetate and β-caryophylline were confirmed using standards (data not shown). The major peak observed at retention time of ∼23.54 was attributed to eugenol while two minor peaks observed at 24.98 and 27.62 belonged to eugenyl acetate and β-caryophylline, respectively.

**Figure 5.**
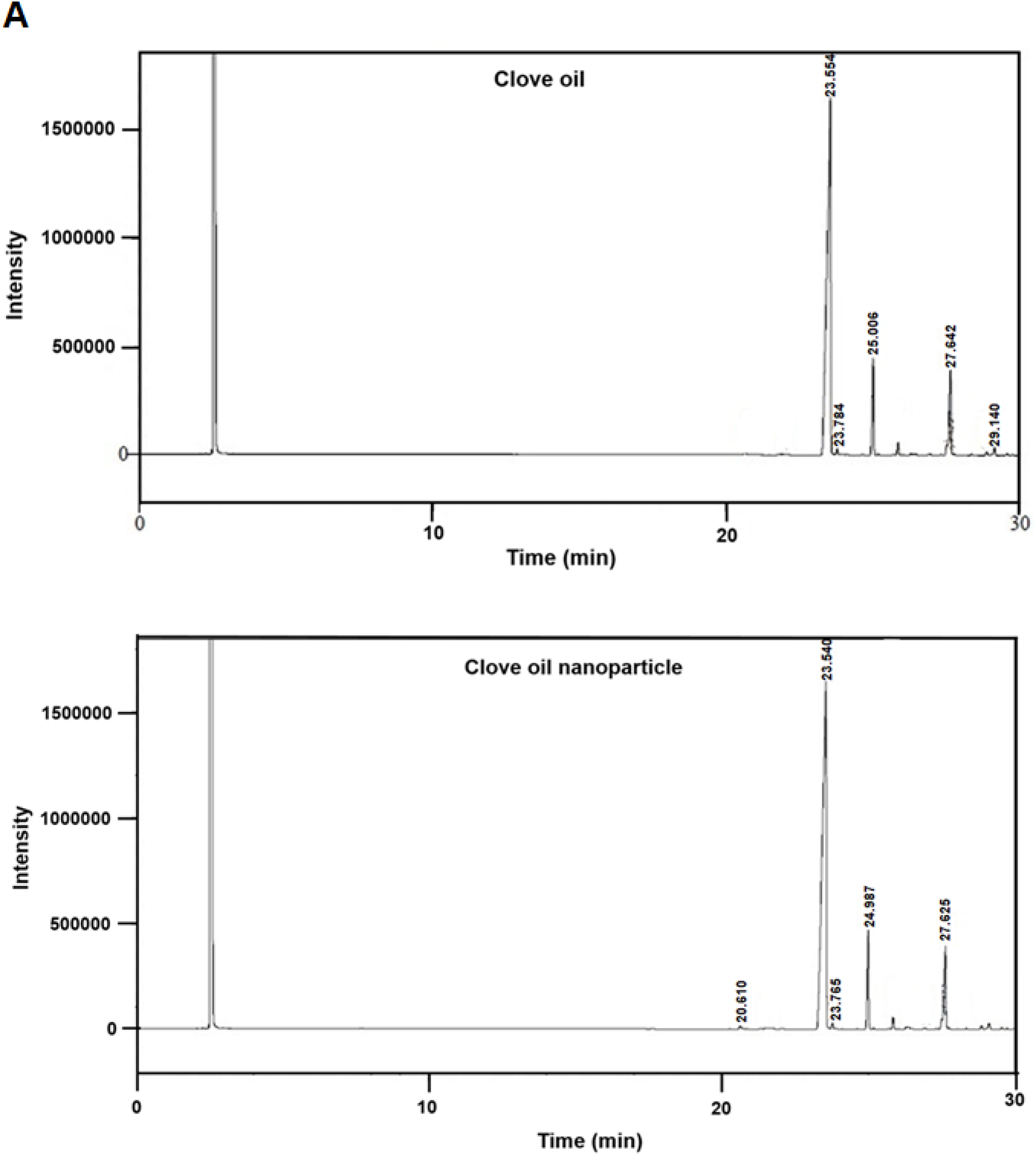
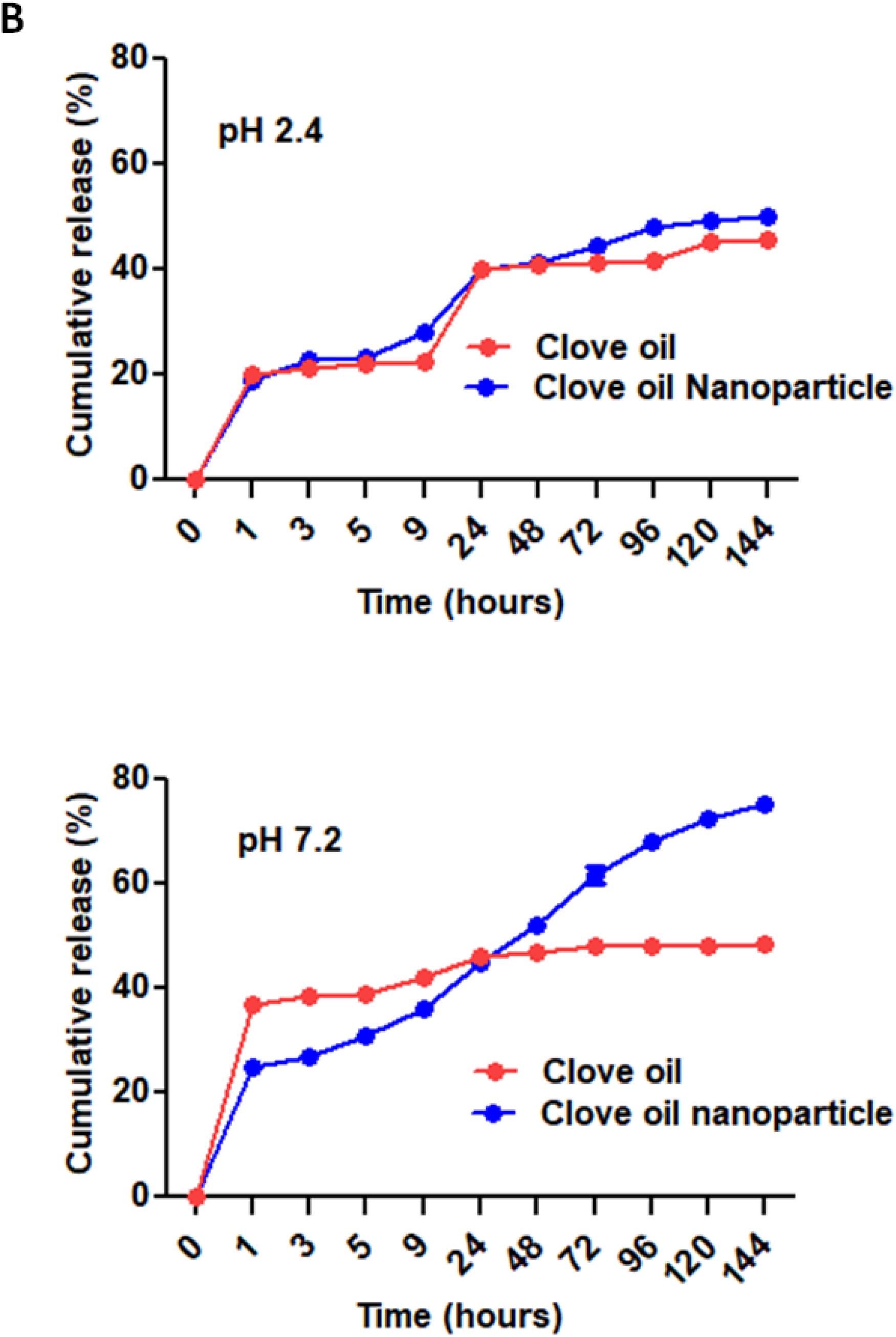
(A) Gas chromatographic profile of Clove oil and Clove oil nanoparticle, (B) *In -vitro* release profile Clove oil of and Clove oil nanoparticle at gastric pH 2.4 and intestinal pH 7.2.

### 2.5. Entrapment efficiency and *in-vitro* release

The clove oil containing eugenol in major proportion and eugenyl acetate, β-caryophylline in small proportion, was dissolved in DCM to prepare a clear solution. The solution was scanned from 250-350 nm in UV-VIS spectrophotometer to identify its λmax and the maximum absorption was obtained at 282 nm. The entrapment efficiency of the spray dried CON samples prepared with Tween 20, 40 and 80 was in the range of 93-98%. This showed the non-ionic surfactant with different HLB did not affect the encapsulation ability substantially in freshly prepared and spray-dried powder. The high entrapment efficiency of CON with Tween 80 was further confirmed using GC spectra data where 98% clove oil retention was calculated. The CON sample (Tween 80) with 98% encapsulation was selected for comparative *in vitro* release study of NC and CON for 144 hrs. The cumulative release of NC at pH 2.4 and 7.2 was 47% and 44%, respectively showing an low solubility at both pH. Further, the profile showed uncontrolled and irregular release. The cumulative percentage release of clove oil from CON was fast in initial 9 hrs. showing 27-35% release followed by a sustained release up-to 49% at pH 2.4 and ∼77% at pH 7.2 (Fig. 5B). The initial burst of release was attributed to the clove oil adsorbed on surface and available at the interface while the sustained release was due to the embedded clove oil. Thus, the nanoencapsulation facilitated controlled release of clove oil. The higher release of clove oil from the nanoparticle at the intestinal pH was due to the higher solubility of maltodetrin at pH 7.4 than in gastric pH 2.4 [31].

### 2.6. Cytotoxicity and hemolytic activity

Understanding the toxicity and hemolytic activity of nanoparticles is an important parameter to evaluate the safety at cellular level and circulation characteristics of nanoencapsulated clove oil. Further, the *in vitro* cytotoxicity of nanoparticles varies with respect to their size and cell type [32, 33]. Hence, a cell viability assay was carried out to investigate the *in vitro* toxicity profile of encapsulated CON on Caco2 cell lines using MTT. As shown in Fig. 6A, CON exhibited greater cell viability when compared to NC. Even at highest concentration of 200 μg/mL, the cell toxicity of CON was 57%, whereas a similar toxicity was observed for NC at the lowest concentration of 20 μg/mL. The hemolytic assays showed that the CON was hem compatible with less than 5 % hemolysis up to10 µg/mL which is below the safe hemolytic ratio for biomaterials according to ISO/TR7406 (Fig. 6B). Further, not more than 10% of hemolysis was observed for all tested concentrations up to 50 µg/mL. However, at higher concentrations, NC showed increased percentage of haemolytic activity when compared to CON. Thus, both the data demonstrates that the nanoencapsulation significantly reduces the cytotoxicity of clove oil.

**Figure 6.**
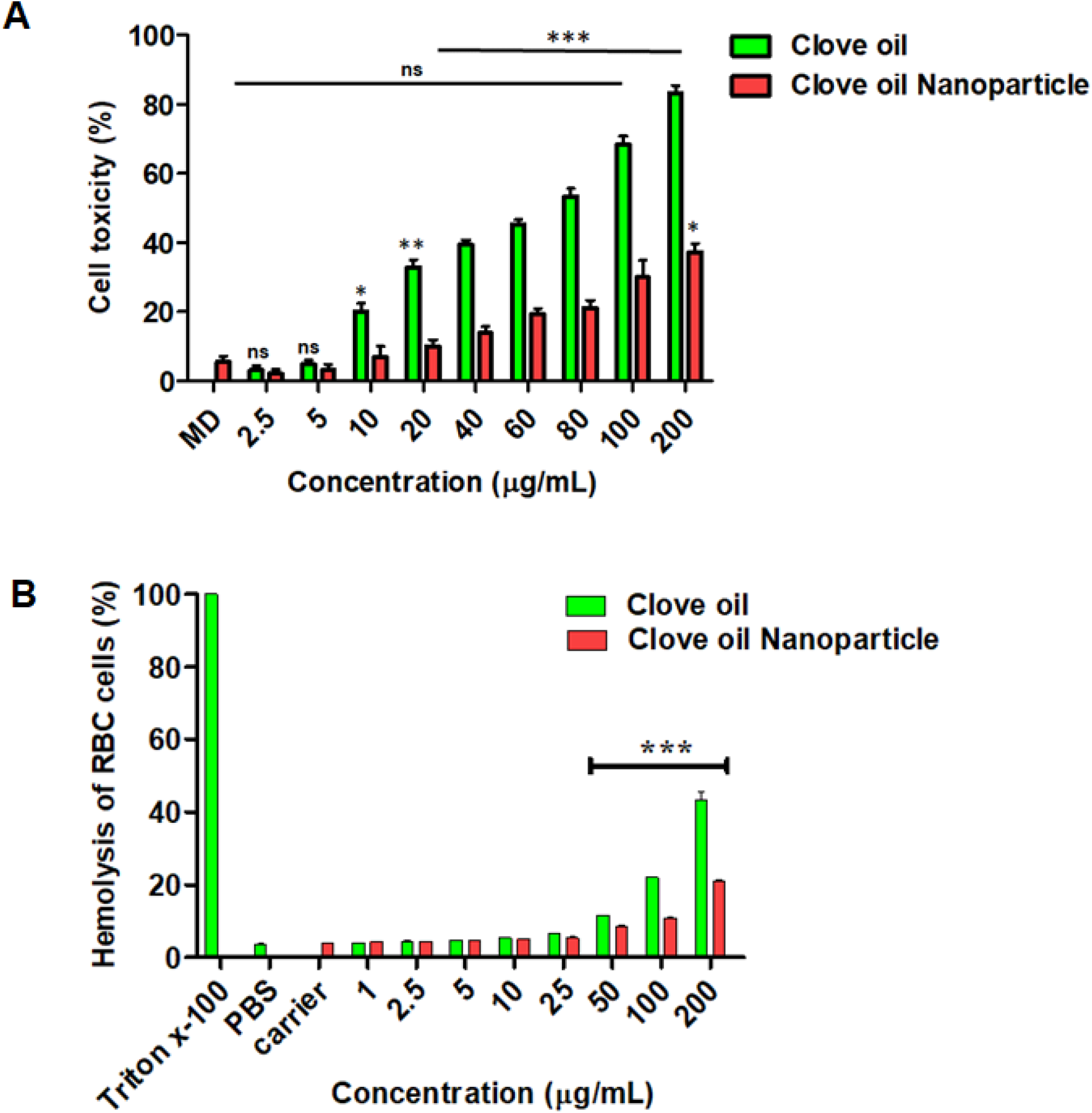
Cytotoxicity and Haemolytic activity. (A) The cell toxicity of Clove oil and Clove oil nanoparticle was measured by MTT assay using human epithelial colorectal adenocarcinoma cells (Caco2). The concentrations of Clove oil and Clove oil nanoparticle used in the experiments were in the range of 2.5 to 200 µg/mL. Maltodextrin was acted as carrier control (B) Percentage of hemolysis of red blood cells after exposure to clove oil and Clove oil nanoparticles at concentrations of 1 µg to 200 µg/mL. the Triton X-100 (1 %) was used haemolytic control. Values are represented as mean SD of experiments done in triplicates. ***p < 0.001, ***p<0.01. ns (not significant) compared to maltodextrin control

### 2.7. Antioxidant potential of CON

Nanoencapsulation is known to enhance antioxidant potential of natural molecules [34, 35]. Hence, free radical-scavenging activity was evaluated by measuring the scavenging activity of the CON on DPPH and ABTS radicals. As shown in Fig. 7A, CON reduced the DPPH radical formation in a dose-dependent manner. The CON showed a maximum inhibition of 80% at a concentration of 100µg/mL, whereas NC at the same concentration exhibited 40% of inhibition. The calculated EC_50_ values of NC and CON were 38.89 and 21.25 μg/mL respectively. ABTS assay is operable over a wide range of pH, is inexpensive, and more rapid than that of DPPH assay and hence the ABTS radical scavenging activity of the CON was investigated [1, 36]. As shown in Fig. 7B, CON was most effective with the EC_50_ of 39.99 μg/mL when compared to NC, which was 72.12 μg/mL. Overall, our results showed that the encapsulated form of the clove oil had a much higher antioxidant activity than the non-encapsulated clove oil.

**Figure 7.**
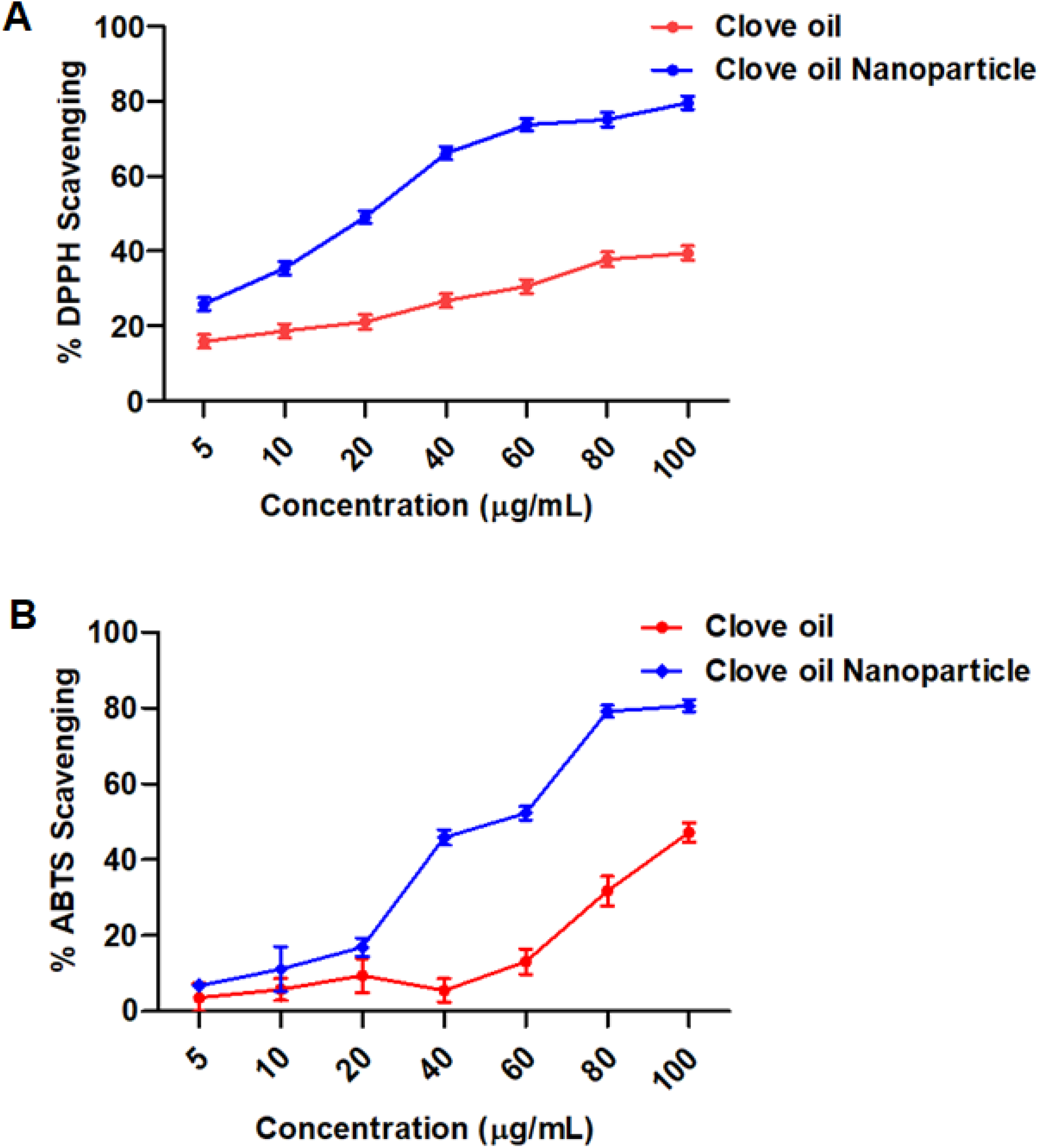
Antioxidant activities of Clove oil and Clove oil nanoparticles (A) DPPH (1, 1-diphenyl picryl hydrazyl) and (B) ABTS ((2,2’-azino-bis (3-ethylbenzothiazoline-6-sulfonic acid)) assays were performed to study the antioxidant activities of Clove oil and Clove oil nanoparticles at concentration ranging from 5-100µg/mL. Quantitation of the results from three independent experiments (n=3) is shown as mean ± SD with statistical significance as *p<0.001 and **p<0.01

## 3. Conclusion

The study highlighted the influence of non-ionic surfactant with different HLB on the stability of clove oil nanoemulsion (CON) and the dried clove oil nanoparticles. Tween 80 having HLB = 15 and the presence of an unsaturated bond at aliphatic chain has resulted in formation of nanoemulsion with small particle size and low ζ. The CON was stable for 8 days while the spray dried clove oil nanoparticles were stable for > 1 month. Thus, masking of pungency, higher water solubility and stability achieved from the study makes CON a potential candidate for drug delivery and to develop nutraceuticals. In addition, nanoencapsulation of clove oil resulted in the controlled release, increased cell viability, blood compatibility and enhanced antioxidant activity compared to native oil. This difference may have a substantial impact on their biological activities and potential health benefits. However, further studies evaluating the *in vivo* toxicity of CON in animal models would enhance its use in biomedical applications.

## 4. Materials and Methods

### 4.1. Materials

Clove oil was obtained from Synthite, Bangalore, India. Maltodextrin (Dextrose Equivalent, DE= 7) was gifted by Rockets Laboratories Pvt. Ltd, Mumbai, India. Tween-20, 40 and 80 were purchased from Loba chemie Pvt. Ltd., Mumbai, India. Sodium dodecyl sulphate (Analytical grade), methanol from Sisco Chemical Laboratory, India, dichloromethane (HPLC grade) from Merck, India, phosphate buffer saline from HiMedia, India, Uranyl acetate, ABTS, ascorbic acid and potassium persulfate, DPPH reagent from Sigma, India were purchased and used for analysis of sample. Cell culture and other related consumables were procured from Nunc, Thermofisher and Sigma. For TEM analysis Copper-coated carbon grids were purchased from Ted Pella, Inc. Triple distilled (TD) water or Milli-Q water was used for all experiments,

### 4.2. Preparation of nanoparticles

Clove oil loaded nanoemulsions (CON) were prepared using oil-in-water emulsification method followed by ultrasonication (Sonics Ultrasonic processor, model no VCX750, USA) for 30 min. The maltodextrin, 4% (w/v) used as a carrier, was homogenized in 100 mL TD water followed by addition of 0.75 % (v/v) clove oil and 60% Tween-20/40/80 (by volume). The percentage of Tween-20/40/80 depicted in this manuscript is, with respect to volume of clove oil. The nanoemulsions were dried using freeze drier (Alpha 2-4 LD plus) at -40^°^C under a vacuum pressure of 0.001 mbar and spray dried (SD, LSD-48, mini Spray Dryer, JISL) at 110^°^C as inlet temperature and 65^°^C as outlet temperature with 1 mL/minute flow rate.

### 4.3 Particle size and zeta potential measurements

The average particle size and zeta potential of CON were measured using a Zetasizer NanoZS (Malvern Instruments, Malvern, UK). Freshly prepared samples were diluted 10 times and used for the measurement. All the measurements were performed in triplicate with 10 runs for each measurement and the results were expressed as mean size ± standard deviation (S.D). The change in particle size, zeta potential and polydispersity index (PDI) of CON were observed for 8 days at room temperature. The particle size data was extracted in excel sheet and the plot is drawn using SigmaPlot.

### 4.4. Emulsion stability

The stability of CON was determined by measuring the extent of the gravitational phase separation processes such as creaming, phase separation and turbidity. Freshly prepared CON samples (10 mL) was stored in a graduated glass test tube at room temperature for 8 days and observed for their stability parameters.

Creaming: 10 mL of nanoemulsion was poured into a transparent measuring glass tube with a stopper. Creaming on the surface of nanoemulsion was measured at time intervals of 0, 2, 4, 6, and 8 days. Creaming index (%) was calculated using the following equation. 1

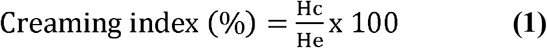

Where, Hc is the height of the cream layer on top and He is the height of total nanoemulsion [37].

Turbidity measurement: 100µl aliquot of CON was diluted with 0.1% sodium dodecyl sulfate (SDS) solution to achieve 0.14-0.16% oil volume fractions. The change in turbidity of samples was measured at 500 nm using UV-Vis spectrophotometer (UV-1800, Shimadzu, UV spectrophotometer) and calculated using the following equation [38].

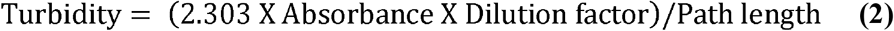

Where, turbidity of emulsion (m^-1^), observed absorbance at 500 nm, path length of cuvette (0.01 m).

### 4.5. Thermal stability

The dissociation temperature of individual reactants and the thermal stability of CON were determined by differential scanning calorimetry (DSC; Perkin Elmer instrument, DSC 800, France). The sample was taken (approximately 3.8 mg) in an aluminium pan and hermetically sealed for the analysis. The samples were heated from 30°C to 295°C with an increase of 5°C/min.

### 4.6. Fourier transform infrared (FTIR) spectroscopy

FTIR spectra of maltodextrin, native clove oil (NC), CON and spray dried powder was recorded using FTIR spectrometer (Perkin-Elmer spectrum one FT-IR spectrometer, USA). The spectra were recorded between 400 cm^-1^ and 4000 cm^-1^ using a high-energy ceramic source and deuterated lanthanum α-alanine doped triglycine sulphate (DLaTGS) detectors.

### 4.7. Scanning electron microscope (SEM) analysis

The spray and freeze dried CON was mounted on carbon-coated specimen holder. FEI ESEM Quanta 200 scanning electron microscope was used to capture their morphologies at an accelerating voltage of 20 kV in high vacuum mode using Everhart-Thornley detector.

### 4.8. Transmission electron microscopy (TEM)

The TEM image of CON and spray dried nanoparticles were captured using Tecnai T-20. The diluted samples were spotted and incubated on the carbon coated copper grid of 400 meshes for 45 seconds. The excessive sample was removed by washing the grids with water. The coated sample was stained using 2% uranyl acetate for 1 minute, the excess of uranyl acetate was removed with a filter paper, and stained samples were observed under TEM at 200 kV.

### 4.9. Entrapment efficiency- Gas chromatography (GC), UV-Vis spectroscopy

Entrapment efficiency of spray dried CON was determined using GC and UV-Vis spectrophotometer. For GC analysis, 1 mL of CON was dissolved in 5 mL of dichloromethane (DCM) and centrifuged at 7,000*g* for 15 minutes at 25°C to remove the adsorbed clove oil. The pellet was collected, dissolved in water and vortexed vigorously. The solution was poured in a separating funnel after mixing with DCM and allowed for phase separation. The process was repeated thrice and the organic phase was collected each time. The DCM extracted clove oil was analysed and compared with NC using gas chromatography (Perkin Elmer GC clarus-580) equipped with a FID and an Elite-5 fused-silica column (30 m, 0.25 mm internal diameter, 0.25 µm film thickness, Schimadzu, Japan). Initial oven temperature was maintained at 40°C for 2 min followed by programmed heating from 40 to 260°C at a rate of 5°C/min, and at 260°C held for 2 min. Injector and detector were set at temperatures of 250°C and 265°C, respectively. The inert carrier gas, nitrogen, was adjusted to a linear velocity of 25.6 cm/sec. Two microliter of NC or clove oil extracted from CON dissolved in DCM was injected into the GC by split mode with a split ratio of 1/20. The standard curve constructed using native clove oil in DCM was used to calculate the entrapment efficiency.

UV-Vis spectrophotometer was used to find the maximum absorption wavelength (λ_max_) of clove oil dissolved in DCM. The sample was scanned from 250 to 350 nm and the maximum absorption was recorded at 282 nm. Hence, the 282 nm was used to record the absorbance of CON taking DCM as blank. The percentage of entrapment efficiency of CON was calculated using a standard curve (R^2^ = 0.9987) and with the following formula

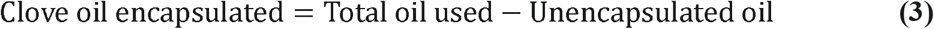

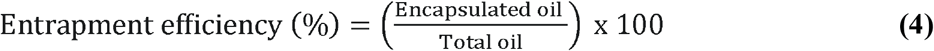

### 4.10. *In vitro* release assay

Two milliliter of CON and NC were taken in equal volume of phosphate buffer solution (PBS; pH 2.4 and 7. 2) and placed in the dialysis bag (MMWCO 12,000, HiMedia, Mumbai, India). Hermetically sealed bags were immersed in 200 mL PBS containing 0.06% Tween-80 and was dialyzed at 37°C with continuous magnetic stirring at 200 rpm [39]. At regular intervals, 1 mL aliquot of the sample was drawn and immediately another 1mL of fresh buffer was added to maintain the constant buffer volume. From the aliquots, the amount of clove oil released from the NC and CON was assessed using UV-Vis spectroscopy at 282 nm. A standard curve was used to determine the concentration of released clove oil and the percentage of release was calculated using following equations,

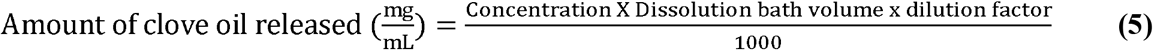

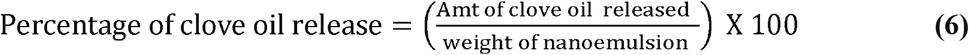

### 4.11. Cell toxicity assay

Caco-2 (human epithelial colorectal adenocarcinoma cell line) cells were purchased from ATCC and cultured in DMEM high glucose media (Sigma). The cells were trypsinized using 0.25% trypsin-EDTA and resuspended in the fresh media. The cells were diluted to 5 × 10^4^ cells/ 100 μL, added to each well of 96-well culture plates and incubated at 37°C for 24 h. The cytotoxic effects of NC and CON was evaluated by treating the cells with various concentrations NC and CON ranging from 2.5-200 µg/mL along with media control for 12 hours at 37°C. Post treatment, 5 mg/mL of MTT (3-(4,5-dimethylthiazol-2-yl)-2,5-diphenyl tetrazolium bromide, Sigma-Aldrich, USA) in DMEM medium was added to each well and incubated at 37°C for another 4□h. The cell viability was assessed by measuring the absorbance at 570 nm wavelength using Epoch 2 microplate spectrophotometer (Agilent technologies. USA).

### 4.12. Hemolysis assay

Hemolytic activity of CON was analyzed by spectrophotometric procedure as described previously with minor modifications [40]. Blood sample (5 mL) was collected from a healthy rat in vials (IAEC No-CFT/IAEC/84/2017) containing EDTA and centrifuged for 5 min at 5000 rpm. Buffy coat was removed and the packed cells were washed three times with 1X PBS (pH 7.4). Further, the pellet was resuspended in 0.5 % saline. Various concentrations of CON (2.5– 200 µg/mL in PBS) was dispensed into 1 mL of erythrocyte suspension individually and incubated at room temperature for 30 min. Triton x 100 (1%) was used as haemolytic control. Post incubation, the samples were centrifuged for 10 min at 5000 rpm and the absorbance of supernatant was measured at 540 nm. The average value was calculated from triplicate readings. The percentage of hemolysis was calculated by dividing absorbance of the sample by positive control absorbance (complete hemolysis) and multiplying with 100 [41].

### 4.13. Antioxidant assay

#### DPPH Assay

The CON and NC were tested for their antioxidant property using 1, 1-diphenylpicrylhydrazyl (DPPH) method [42]. The CON solution (1 mg/mL) was diluted to final concentration of 5, 10, 20, 40, 60, 80 and 100 µg/mL in water. 300 µL of methanolic DPPH solution (1 mL, 0.3 mmol) was added to CON and NC, incubated at room temperature for 30 min in dark and the absorbance was measured at 517 nm in UV VIS-1800 spectrophotometer (Epoch™ 2 Microplate Spectrophotometer) [43]. The scavenging activity was calculated by following formula:

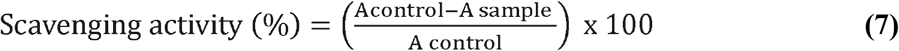

Where, A_control_ and A_sample_ are absorbance of control (DPPH solution) and different samples, respectively. The methanolic DPPH solution (1 mL, 0.3 mmol) was used as control. NC and CON with indicated concentrations were taken to calculate EC_50_ values (Log concentration on the x-axis and their relative percentage inhibition on the y-axis). The R^2^ values were 0.9564 and 0.9934 for NC and CON, respectively. All the experiments were conducted in triplicates.

#### ABTS method

ABTS radical scavenging activity was determined according to the method described by Sridhar *et al* [44] with some modifications. The ABTS stock solution was prepared by adding equal quantities of 7□mM ABTS aqueous solution and 2.45□mM aqueous solutions of potassium persulfate and incubating for 12–16□h at room temperature in dark. 1□mL of this solution was mixed with various concentrations (5-100□µg/mL) of ascorbic acid (used as standard), sample (5, 10, 20, 40, 80 and 100 µg/mL) and incubated at room temperature for 10□min in the dark. The 1□mL of ABTS solution mixed with 0.5□mL of double distilled water was used as control. The absorbance was recorded at 734 nm and the percentage of scavenging activity was calculated using the following equation.

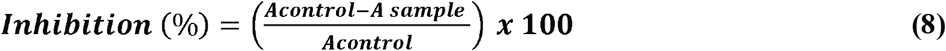

All the experiments were conducted in triplicates. The calibration curves and the EC_50_ values were calculated as mentioned before. The R^2^ values were 0.9563 and 0.9638 for NC and CON, respectively.

## Acknowledgements

The authors are grateful to Director, CSIR-CFTRI, Mysuru, for his constant support. The authors are also thankful to Head & staff, SFS, DMN and CIFS, CSIR-CFTRI, Mysuru for their support. Pramod G Nagaraju acknowledges the fellowship by Department of Science and Technology (DST), New Delhi, India. The Authors thank Dr. Subash Chandra Bose Chinnathambi, CSIR-National Chemical laboratory, Pune for his help in TEM studies

## Summary

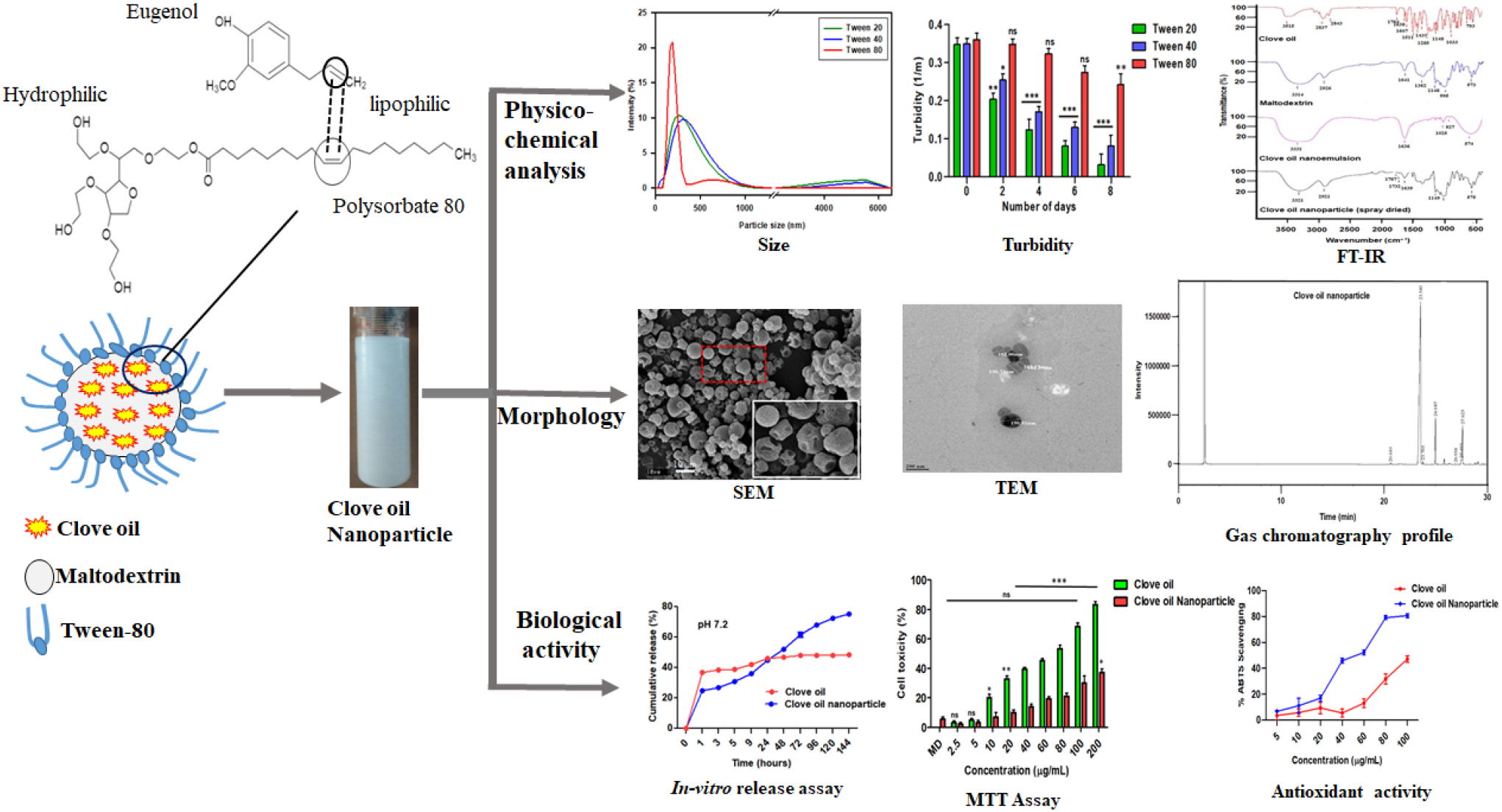

